# Proteomics of osmoregulatory responses in threespine stickleback gills

**DOI:** 10.1101/2020.03.11.987834

**Authors:** Johnathon Li, Dietmar Kültz

## Abstract

The gill proteome of threespine sticklebacks (*Gasterosteus aculeatus*) differs greatly in populations that inhabit diverse environments characterized by different temperature, salinity, food availability, parasites, and other parameters. To assess the contribution of a specific environmental parameter to such differences it is necessary to isolate its effects from those of other parameters. In this study the effect of environmental salinity on the gill proteome of *G. aculeatus* was isolated in controlled mesocosm experiments. Salinity-dependent changes in the gill proteome were analyzed by LC/MSMS data-independent acquisition (DIA) and Skyline. Relative abundances of 1691 proteins representing the molecular phenotype of stickleback gills were quantified using previously developed MSMS spectral and assay libraries in combination with DIA quantitative proteomics. General stress responses were distinguished from osmoregulatory protein abundance changes by their consistent occurrence during both hypo- and hyper-osmotic salinity stress in six separate mesocosm experiments. If the abundance of a protein was consistently regulated in opposite directions by hyper- versus hypo-osmotic salinity stress, then it was considered an osmoregulatory protein. In contrast, if protein abundance was consistently increased irrespective of whether salinity was increased or decreased, then it was considered a general stress response protein. KEGG pathway analysis revealed that the salivary secretion, inositol phosphate metabolism, valine, leucine and isoleucine degradation, citrate cycle, oxidative phosphorylation, and corresponding endocrine and extracellular signaling pathways contain most of the osmoregulatory gill proteins whose abundance is directly proportional to environmental salinity. Most proteins that were inversely correlated with salinity map to KEGG pathways that represent proteostasis, immunity, and related intracellular signaling processes. General stress response proteins represent fatty and amino acid degradation, purine metabolism, focal adhesion, mRNA surveillance, phagosome, endocytosis, and associated intracellular signaling KEGG pathways. These results demonstrate that *G. aculeatus* responds to salinity changes by adjusting osmoregulatory mechanisms that are distinct from transient general stress responses to control compatible osmolyte synthesis, transepithelial ion transport, and oxidative energy metabolism. Furthermore, this study establishes salinity as a key factor for causing the regulation of numerous proteins and KEGG pathways with established functions in proteostasis, immunity, and tissue remodeling. We conclude that the corresponding osmoregulatory gill proteins and KEGG pathways represent molecular phenotypes that promote transepithelial ion transport, cellular osmoregulation, and gill epithelial remodeling to adjust gill function to environmental salinity.

## Introduction

Three-spined sticklebacks (*Gasterosteus aculeatus*) have a ubiquitous geographical distribution range spanning almost the entire Northern hemisphere. Ancestrally marine populations of this species have repeatedly colonized many brackish and freshwater (FW) habitats throughout Europe, Asia, and North America (Paepke 1996). Marine, FW and brackish/ anadromous populations have different morphology regarding body armor (lateral plating and pelvic girdle robustness), size, and head-to-body ratio (Svanbäck and Schluter 2012). Low-plated fish are small and have a large head-to-body ratio (*G. aculeatus leiurus*). They are confined to FW and brackish water that does not exceed isosmolality (McKinnon and Rundle 2002). Fully plated fish are much larger and have a smaller head-to-body ratio (*G. aculeatus trachurus*). They are generally marine or anadromous although some resident FW populations having this morphotype do exist. An intermediate morphotype (partially plated *G. aculeatus semiarmatus*) is also common in population mixing zones such as estuaries (Paepke 1996). In addition to different morphotypes, this species is also characterized by different ecotypes: resident marine, resident FW, and anadromous populations. In general, a low-plated morphotype corresponds to a FW ecotype and a fully plated ecotype corresponds to a resident marine or anadromous ecotype. However, atypical combinations of morpho- and eco-types have also been observed (Paepke 1996). Moreover, different ecotypes also show different behavioral and physiological phenotypes such as migratory behavior, metabolic rate, swimming endurance, and osmoregulation (Kitano et al. 2012; Divino et al. 2016). These phenotypic differences between ecotypes indicate strong selective pressures that promote adaptive processes conferring significant fitness advantages in specific habitats.

While several studies have compared mRNA levels in sticklebacks exposed to different environments using transcriptomics, very few have examined environmental effects on any stickleback tissue using experimental proteomics. We have recently shown that the gill proteome of North American *G. aculeatus* is characterized by large differences between populations that inhabit the northern and southern edges of the distribution range and represent different eco- and morpho-types (Li et al. 2018). The gill was chosen as the target tissue because the branchial proteome is responsible for osmoregulatory, acid-base regulation, nitrogen excretion, and respiratory functions that are essential for growth and survival in marine and FW habitats and for swimming activity and aerobic capacity (Evans and Claiborne 2009). The teleost gill is mostly made up of primary filaments and parallel sheets of secondary lamella where intense blood circulation enables efficient gas exchange, ion transport, pH regulation, and nitrogenous waste excretion. These epithelial structures are supported by cartilaginous tissue that forms the gill arches. Fish gills represent an excellent target tissue for analyzing environmental effects on fish because they are directly exposed to the external medium and responsible for many critical physiological processes such as water and oxygen exchange, osmoregulation, acid-base balance, and nitrogen excretion. In addition, gill epithelium sensitively reflects immune responses of fish and represents a good bioindicator tissue target for health assessment (Austbø et al. 2014).

Because field populations experience heterogeneous environments, differences in any individual or combination of environmental parameters in addition to genetic differences may account for their altered gill proteomes. To isolate the influence of a particular environmental variable on the gill proteome it is necessary to perform mesocosm experiments under controlled conditions using appropriate controls (Somero 2012). Moreover, it is necessary to distinguish non-specific responses to environmental change/ stress that may occur during acclimation to particular environmental parameters from parameter-specific/ directional responses (Kültz 2020). The present study isolates salinity-specific effects on the *G. aculeatus* gill proteome from those of other environmental parameters and genetic differences that distinguish eco- and morpho-types.

## Materials and Methods

### Mesocosm experiments

Juvenile threespine sticklebacks (*Gasterosteus aculeatus*) were collected from Bodega Harbor adjacent to Spud Point Marina in Bodega Harbor (BH, salinity = 34 g/kg) and Lake Solano (LS, salinity = 0.4 g/kg) in August 2012 and June 2014 in accordance with California Department of Fish and Wildlife scientific collecting permits SCP11108 and SCP12637. They were raised in the laboratory over a period of four months. All fish were kept under identical conditions during this four-month pre-acclimation period. Six salinity acclimation experiments were performed. BH fish collected in August 2012 were gradually acclimated from 34 g/kg to 1 g/kg at a rate of 3 g/kg per day over a period of 11 days and kept at the final salinity for an additional 24 h (experiment BH1.1). This experiment included parallel handling controls that were kept at 34 g/kg. Experiment BH67.1 consisted of gradual acclimation of BH fish collected in August 2012 from 34 g/kg to 67 g/kg with parallel handling controls kept at 34 g/kg. These two experiments were repeated with a different cohort of fish collected in June 2014 to account for effects of different generations and yearly variations in climate on salinity-induced proteome variation (experiments BH1.2 and BH67.2). In addition, LS fish collected in August 2014 were acclimated at a rate of 3 g/kg per day from 1 g/kg to 34 g/kg (experiment LS34) and from 1 g/kg to 67 g/kg (experiment LS67). Handling controls (1 g/kg → 1 g/kg) were also included in these experiments. Using both BH (marine *G. aculeatus trachurus*) and LS (FW *G. aculeatus leiurus*) populations for these mesocosm experiments supports identification of osmoregulatory and general stress response proteins whose salinity-regulation is conserved in different eco- and morphotypes of this species. All experiments were approved by the University of California Davis Institutional Animal Care and Use Committee (IACUC number 18010, AAALAC number A3433-01).

### Sample preparation and peptide separation for proteomics

Following euthanasia of fish, gills were dissected, snap-frozen in liquid nitrogen and stored immediately at −80°C until further processing. Protein extraction, protein assay, and in solution trypsin digestion were performed as previously reported (Kültz et al. 2013). Briefly, proteins were digested using sepharose bead-immobilized porcine trypsin (Promega) at a ratio of 50 : 1 (sample peptide versus trypsin concentration) in a rotator overnight at 35°C. Tryptic peptides from each sample (200 ng total) were injected with a nanoAcquity sample manager (Waters, Milford, MA), trapped for 1 min at 15 μL/min on a Symmetry trap column (Waters 186003514), and separated on a 1.7μm particle size BEH C18 column (250mm × 75μm, Waters 186003545) by reversed phase liquid chromatography using a nanoAcquity binary solvent manager (Waters). A 125 min linear gradient ranging from 3% to 35% acetonitrile (ACN) was used. Peptides were directly eluted (online) into a UHR-qTOF mass spectrometer (ImpactHD, Bruker) using a pico-emitter tip (New Objective FS360–20-10-D-20, Woburn, MA). Batch-processing of samples was controlled with Hystar 4.1 (Bruker) and a quality control standard (68 fmol BSA peptide mix) was included on a weekly basis to monitor instrument performance.

### Data-independent acquisition (DIA) and quantitation of MSMS spectra

Eluted peptides were ionized by nano-electrospray ionization (nESI) followed by separation in a quadrupole mass analyzer, collision-induced fragmentation (CID) and additional separation of fragment ions in a time-of flight mass analyzer. A previously constructed and validated DIA assay library for *G. aculeatus* gills was used to quantify the same set of transitions, peptides, and proteins in each sample (Li et al. 2018). The mass range for DIA was set to 390 – 1015 m/z at 25 Hz scan rate with an isolation width of 10 m/z (1 m/z overlap, 2.5 sec scan interval). Quantitative analyses and visualization of DIA data was performed using Skyline 19.0. At least five (generally six) transition peaks were detected and scored using the mProphet algorithm integrated into Skyline. The mass error threshold for transitions was 25 ppm and the resolving power was 30,000. Randomly scrambled decoys were used to calculate mProphet Q values. MSstats 3.1, which is an R package specifically designed for the statistical analysis of DIA quantitative proteomics data that is integrated into Skyline, was used for power analysis to calculate the fold-change (FC) cutoff that is appropriate for each experiment. MSstats 3.1 was also used to determine statistically significant differences between treatments and corresponding handling controls. The cutoff for multiple testing (Benjamini-Hochberg) adjusted p-values was set to p<0.05. The normalization method for MSstats analyses was to equalize medians at a minimum confidence interval of 95% and with protein quantity as the scope for each analysis. The summary method used for MSstats analyses was Tukey’s median polish to give equal weighting to each transition and each peptide of a given protein and the minimal mProphet detection Q value for peak quality was set to 0.05.

### KEGG pathway analysis

The consistency of salinity-dependent regulatory patterns of gill proteins in all six mesocosm experiments was determined using Excel (Microsoft) and visualized with Venny 2.1 (Oliveros 2015). Only those DIA assay library proteins that were statistically significantly regulated by salinity in at least one experiment were considered for pattern analysis. Four regulation patterns suitable for discerning osmoregulatory from general stress responses were evaluated as follows: 1) osmoregulatory proteins that were consistently upregulated when salinity increased and downregulated when salinity decreased; 2) osmoregulatory proteins that were consistently inversely correlated to a salinity change (opposite pattern as that of pattern 1); 3) general stress response proteins that were always upregulated, independent of the direction of salinity stress; 4) proteins that were always downregulated, independent of the direction of salinity stress. The sets of proteins matching these four regulation patterns were matched to KEGG pathways using KEGG mapper after annotating the entire *G. aculeatus* gill assay library with KEGG orthology (KO) identifiers using GhostKOALA (Kanehisa et al. 2016). Quantitative heat maps of KEGG pathways reflecting protein abundance FC values for each experiment were generated using the KEGG mapper color pathway tool (Kanehisa and Sato 2020).

## Results

### Visualization of high-dimensional DIA quantitative proteomics data

The DIA assay library used for label-free quantitative proteomics of *G. aculeatus* gill samples has been constructed and validated in a previous study (Li et al. 2018). It contains 1691 proteins that are represented by 4746 peptides, 4879 precursors (specific peptide charge states), and 26,123 transitions (MSMS fragment ions that are being monitored). Thus, for each sample a very complex, high-dimensional data set is being generated, which increases further in complexity when multiple samples are being compared within an experiment and for multiple experiments. To properly capture, analyze, visualize, and comprehend the high-dimensional nature of these data sets we utilized Skyline, which has been specifically developed for DIA quantitative proteomics and other targeted LC-MSMS experiments (Pino et al. 2017). All data and metadata associated with a sample and experiment are accessible through Skyline and the associated open-access repository Panorama-Public (Sharma et al. 2014). For each protein, quantitative data are visually accessible in the form of extracted ion chromatograms (XICs) for each transition (Figure 1). In addition, metadata, library spectra, and peak quality criteria such as the intensity pattern of peptide fragment ions relative to each other (dotp value) are displayed. An example is shown for Na^+^/K^+^/2Cl^−^ cotransporter NKCC (Uniprot G3Q307) quantitation in experiment BH1.1, in which marine Bodega Harbor fish were transferred from 34 g/kg to 1 g/kg (Figure 1). This example illustrates that even for low abundance proteins with low peptide ion intensities the corresponding peaks can be reliably detected. Moreover, it is evident from this example that the ionization efficiency of unique peptides derived from the same protein can differ greatly, e.g. an order of magnitude between peptides ELPPILLVR^++^ and ATSIQNSPAMQK^++^. Nevertheless, ionization efficiency differences among peptides derived from a single protein do not affect relative quantitation of protein abundance since Skyline and MSstats only directly compare transitions derived from the same peptide. Figure 1 illustrates that sample-specific differences in the relative abundance of all transitions for the three different peptides are very consistent. The corresponding data and metadata for all proteins represented in the DIA assay library and for all samples and experiments analyzed in this study are available at the Panorama-Public targeted quantitative proteomics repository at the following addresses: https://panoramaweb.org/b9QCDP.url. All LC-MSMS raw data have been deposited at ProteomeXchange under accession number PXD017662 (http://proteomecentral.proteomexchange.org/cgi/GetDataset?ID=PXD017662).

**Figure 1:**
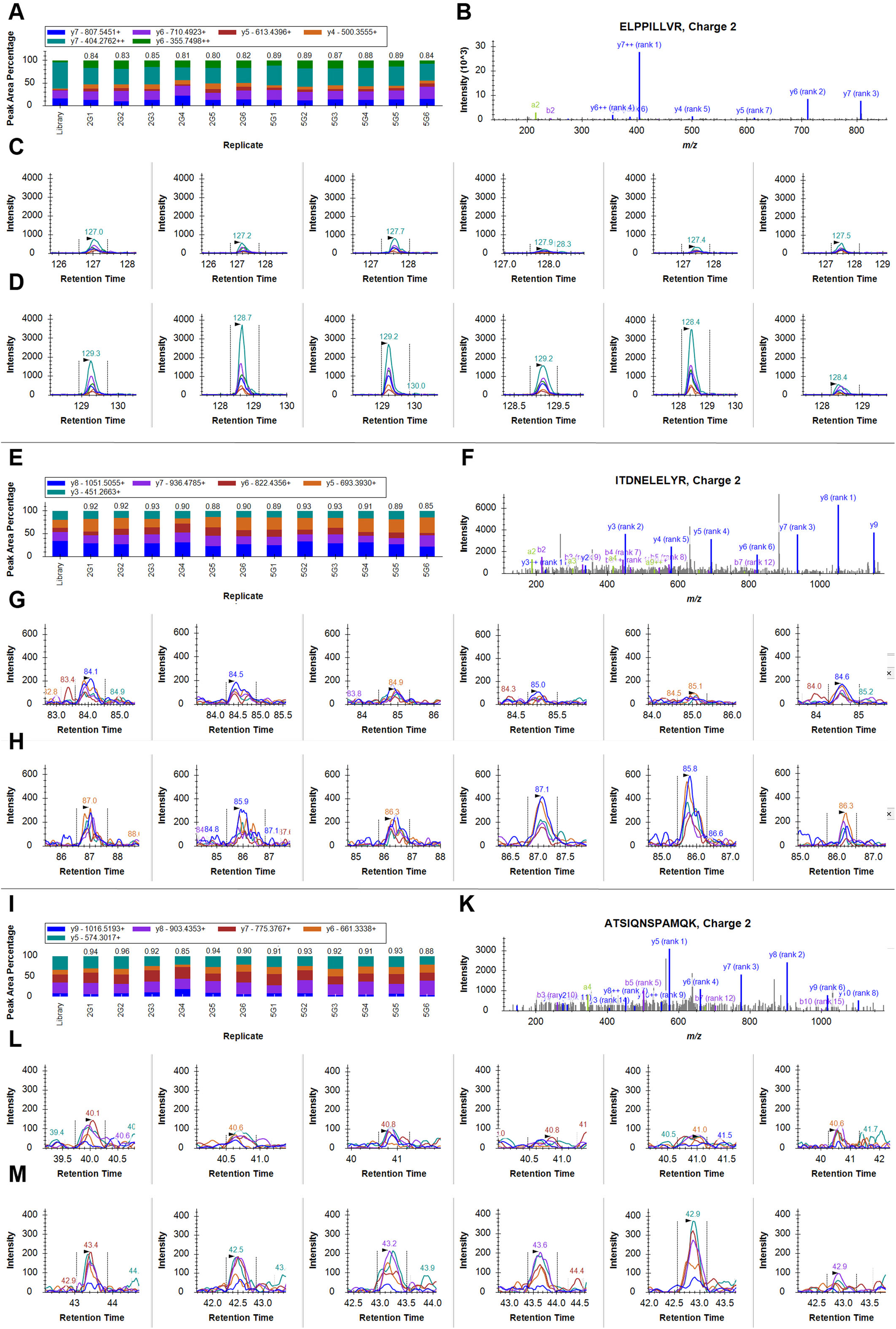
DIA quantitation of the Na^+^/K^+^/2Cl^−^ cotransporter NKCC (Uniprot G3Q307) for all twelve biological replicates of experiment LS67 based on three unique peptides (ELPPILLVR^++^ panels A-D, ITDNELELYR^++^ panels E-H, ATSIQNSPAMQK^++^ panels I-M). In each sample, the intensity pattern of the corresponding transitions, mass/charge (m/z), charge, and retention time is matched to that in the reference library spectrum and assigned a dotp value, which is depicted above the color-coded fragment ion intensity pattern for each sample (panels A, E, and I). The first six samples represent Bodega Harbor fish acclimated to 1 g/kg while the remaining six samples represent handling controls kept at 34 g/kg. The dotp value ranges between 0 (no match) and 1 (perfect match). The legend at the top indicates the color-code used for individual transitions, which match specific fragment ions of the corresponding reference library spectra shown in panels B, F, and K. Individual transitions monitored for each peptide are distinguished by the same color-code in panels C, D, G, H, L, and M. Transitions for six biological replicates acclimated from 34 g/kg to 1 g/kg are shown in panels C, G, and L. Transitions for six control fish maintained at 34 g/kg to 1 g/kg are shown in panels D, H, and M.

### Magnitude of salinity effects on the gill proteome of marine and FW sticklebacks

Gradual acclimation of BH sticklebacks from 34 to 1 g/kg (experiment BH1.1) resulted in a significant abundance increase of 102 proteins and decrease of 111 proteins at a fold change (FC) cutoff of 1.9 (n = 6), which was calculated as the FC threshold using the variation inherent in the data for this experiment (Figure 2A,B). BH sticklebacks that were gradually acclimated from 34 to 67 g/kg (experiment BH67.1) showed a significant abundance increase of 83 proteins and decrease of 65 proteins at an FC threshold of 1.9 (n = 6, Figure 2C,D). These two experiments were repeated with sticklebacks that had been collected two years later yielding data sets with greater variances that necessitated an FC threshold increase to 2.9 (experiment BH1.2, n = 8, Figure 3A) and 2.0 (experiment BH67.2, n = 8, Figure 3C) to maintain identical statistical power. In contrast to experiment BH1.1, none of the significantly regulated proteins exceeded this higher FC threshold of 2.9 in the repeat experiment BH1.2 (acclimation from 34 to 1 g/kg, Figure 3B). However, a significant abundance increase of 105 proteins and a decrease of 85 proteins was observed in experiment BH67.2, which represented a repeat of the acclimation from 34 to 67 g/kg performed two years earlier in experiment BH67.1 (Figure 3D). These results demonstrate that FC thresholds and intrinsic noise, i.e. differences in confidence intervals associated with the quantity of each protein, influence the number of proteins that can be identified as significantly altered in abundance by a change in salinity. To identify the most highly conserved fraction of the gill proteome that is regulated by environmental salinity in this species another set of experiments was performed on a resident FW population from Lake Solano. This population represents a different ecotype (land-locked FW) and morphotype (*leiurus*) than the BH population (resident marine *trachurus*). Identification of gill proteome abundance changes that occur in both populations independent of ecotype and morphotype enable assessment of general trends for salinity effects on this species. Gradual acclimation of LS fish from 1 to 34 g/kg (experiment LS1) resulted in a significant abundance increase of 9 proteins and decrease of 4 proteins at an FC threshold of 2.0 (n =12, Figure 4A,B). LS fish that were gradually acclimated from 1 to 67 g/kg (experiment LS67) showed a significant abundance increase of 76 proteins and decrease of 80 proteins at an FC threshold of 2.5 (n = 12, Figure 4C,D). Quantitative data for all proteins and all experiments are summarized in table S1.

**Figure 2:**
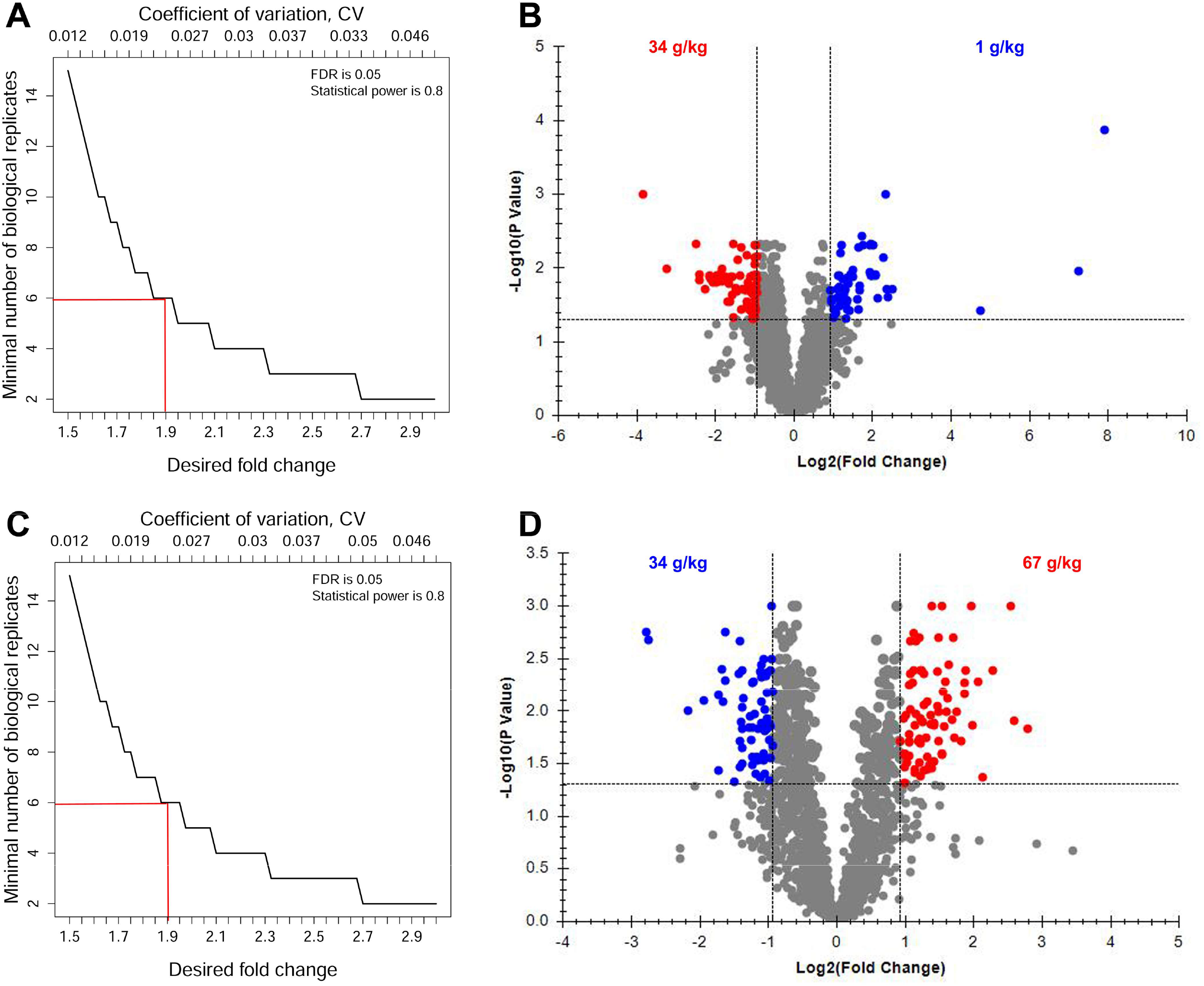
Effect of environmental salinity on the proteome of marine Bodega Harbor sticklebacks collected in summer 2012. **A)** MSstats power analysis that considers the fold change (FC) confidence intervals associated with the abundance of each protein in every sample of experiment BH1.1 (acclimation of fish from 34 g/kg to 1 g/kg). **B**) Volcano plot depicting the relationship between FC and multiple-testing adjusted P value (adj.P) for experiment BH1.1. **C)** MSstats power analysis for experiment BH67.1 (acclimation of fish from 34 g/kg to 67 g/kg). **D**) Volcano plot depicting the relationship between FC and adj.P for experiment BH67.1. Proteins whose abundance is significantly altered by salinity are indicated in red (positively correlated with salinity) and blue (negatively correlated with salinity). Note that FC is expressed as log2(FC) on the abscissa and adj.P as Log10(adj.P) on the ordinate of volcano plots.

**Figure 3:**
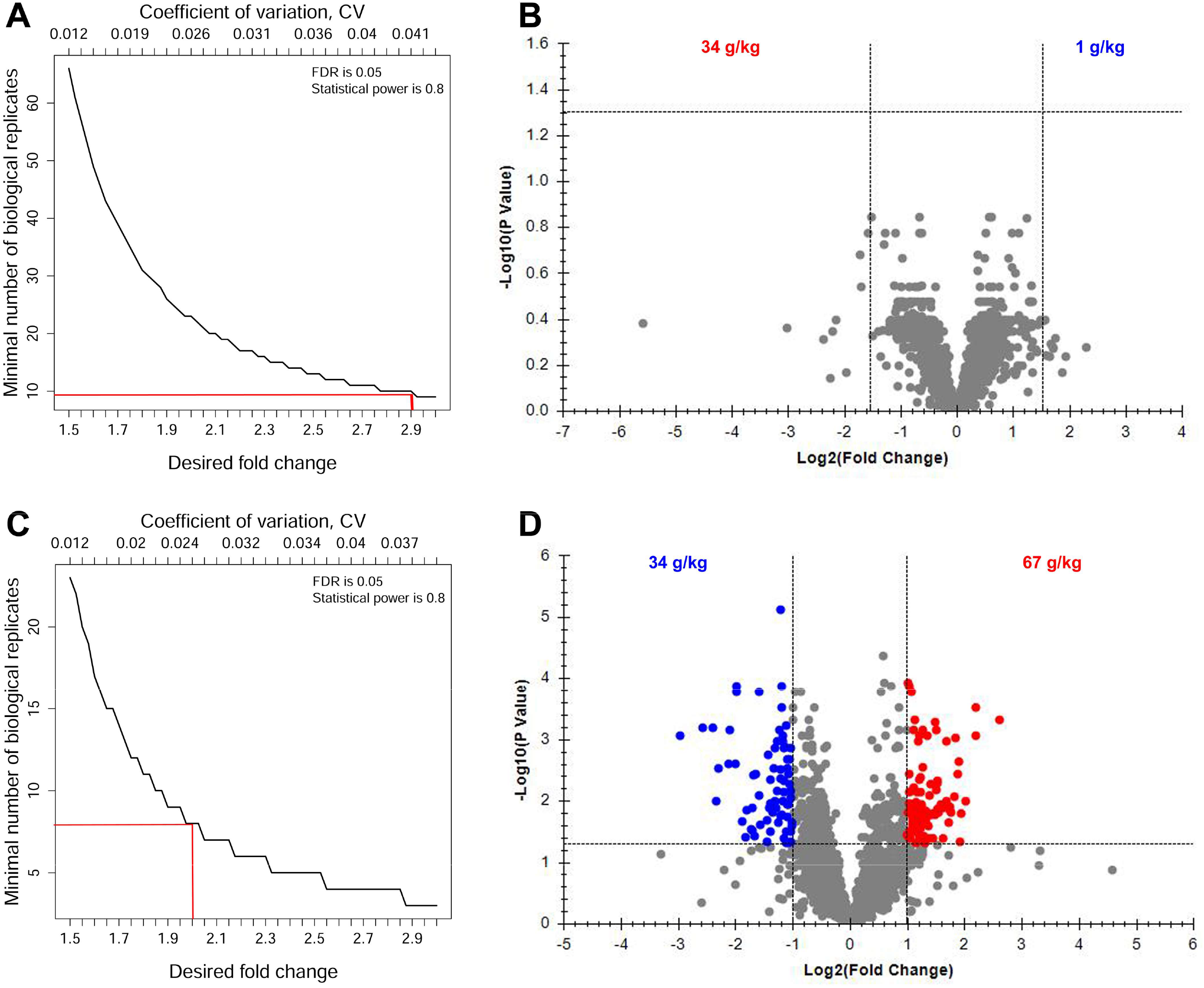
Effect of environmental salinity on the proteome of marine Bodega Harbor sticklebacks collected in summer 2014. **A)** MSstats power analysis that considers the fold change (FC) confidence intervals associated with the abundance of each protein in every sample of experiment BH1.2 (acclimation of fish from 34 g/kg to 1 g/kg). **B**) Volcano plot depicting the relationship between FC and multiple-testing adjusted P value (adj.P) for experiment BH1.2. **C)** MSstats power analysis for experiment BH67.2 (acclimation of fish from 34 g/kg to 67 g/kg). **D**) Volcano plot depicting the relationship between FC and adj.P for experiment BH67.2. Proteins whose abundance is significantly altered by salinity are indicated in red (positively correlated with salinity) and blue (negatively correlated with salinity). Note that FC is expressed as log2(FC) on the abscissa and adj.P as Log10(adj.P) on the ordinate of volcano plots.

**Figure 4:**
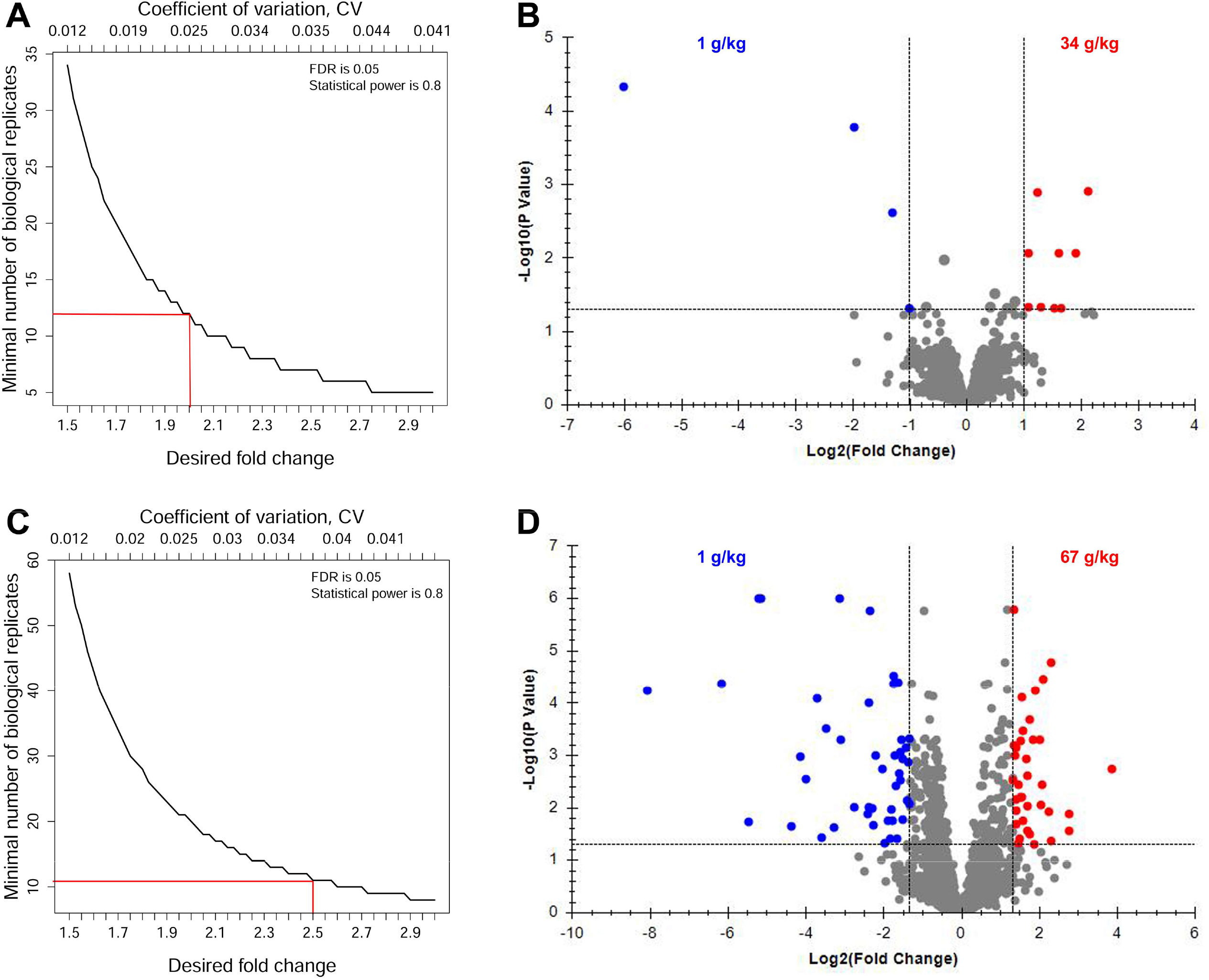
Effect of environmental salinity on the proteome of freshwater Lake Solano sticklebacks collected in summer 2014. **A)** MSstats power analysis that considers the fold change (FC) confidence intervals associated with the abundance of each protein in every sample of experiment LS34 (acclimation of fish from 1 g/kg to 34 g/kg). **B**) Volcano plot depicting the relationship between FC and multiple-testing adjusted P value (adj.P) for experiment LS34 **C)** MSstats power analysis for experiment LS67 (acclimation of fish from 1 g/kg to 67 g/kg). **D**) Volcano plot depicting the relationship between FC and adj.P for experiment LS67. Proteins whose abundance is significantly altered by salinity are indicated in red (positively correlated with salinity) and blue (negatively correlated with salinity). Note that FC is expressed as log2(FC) on the abscissa and adj.P as log10(adj.P) on the ordinate of volcano plots.

### Distinction of osmoregulatory versus general stress response proteins

All proteins that were significantly altered in their abundance in at least one experiment were evaluated for the consistency of their regulatory pattern in all six experiments. The abundance of 48 proteins was directly proportional to the environmental salinity in all six experiments, i.e. they were up-regulated when salinity increased and down-regulated when salinity decreased (Figure 5a, Table S1). These proteins were considered osmoregulatory proteins. They included well-known ion transporters such as the Na^+^/K^+^-ATPase (NKA) alpha (G3PMS4) and beta (G3PRG7) subunits and Na^+^/K^+^/2Cl^−^ cotransporter (G3Q307), and the compatible osmolyte synthesizing enzyme inositol monophosphatase (IMPA1.1, G3NMA6). Forty-six of these proteins mapped to KEGG orthology (KO) identifiers and were, thus, amenable for KEGG pathway analysis. The only two proteins in that group that did not map to any KO identifiers were inter-alpha-trypsin inhibitor heavy chain 3 (G3N9S1) and uridine nucleosidase 1 (G3PPQ2). Fifty-four proteins that showed the opposite pattern of regulation, i.e. their abundance changed inversely proportional to salinity (Figure 5b, Table S1). Nine of these 54 proteins did not match any KO identifier but several of these nine, notably two isoforms of Golgi-associated plant pathogenesis protein (G3NGY5, G3NGY8) and interferon-induced protein 44 (G3NYZ2) had immunity-related functions. Forty proteins consistently increased in abundance in all six experiments, independent of whether salinity was increased or decreased (Figure 5c, Table S1). These proteins were classified as general stress response proteins. The three of them that did not map to any KO identifier were translationally-controlled tumor protein homolog 1 (G3NUU3), actinoporin-like protein (G3P0X6), and grancalcin (G3PL83). Sixty-five proteins were consistently down-regulated in all experiments, independent of whether salinity increased or decreased (Figure 5d, Table S1). These proteins were general stress-repressed proteins. Four of these 65 proteins did not map to any KO identifier, including sarcolemmal membrane-associated protein (G3NE74), tumor protein D52 (G3NGP2), LIM and SH3 domain protein LASP1 (G3NH50) and secretagogin (G3NUX2).

**Figure 5:**
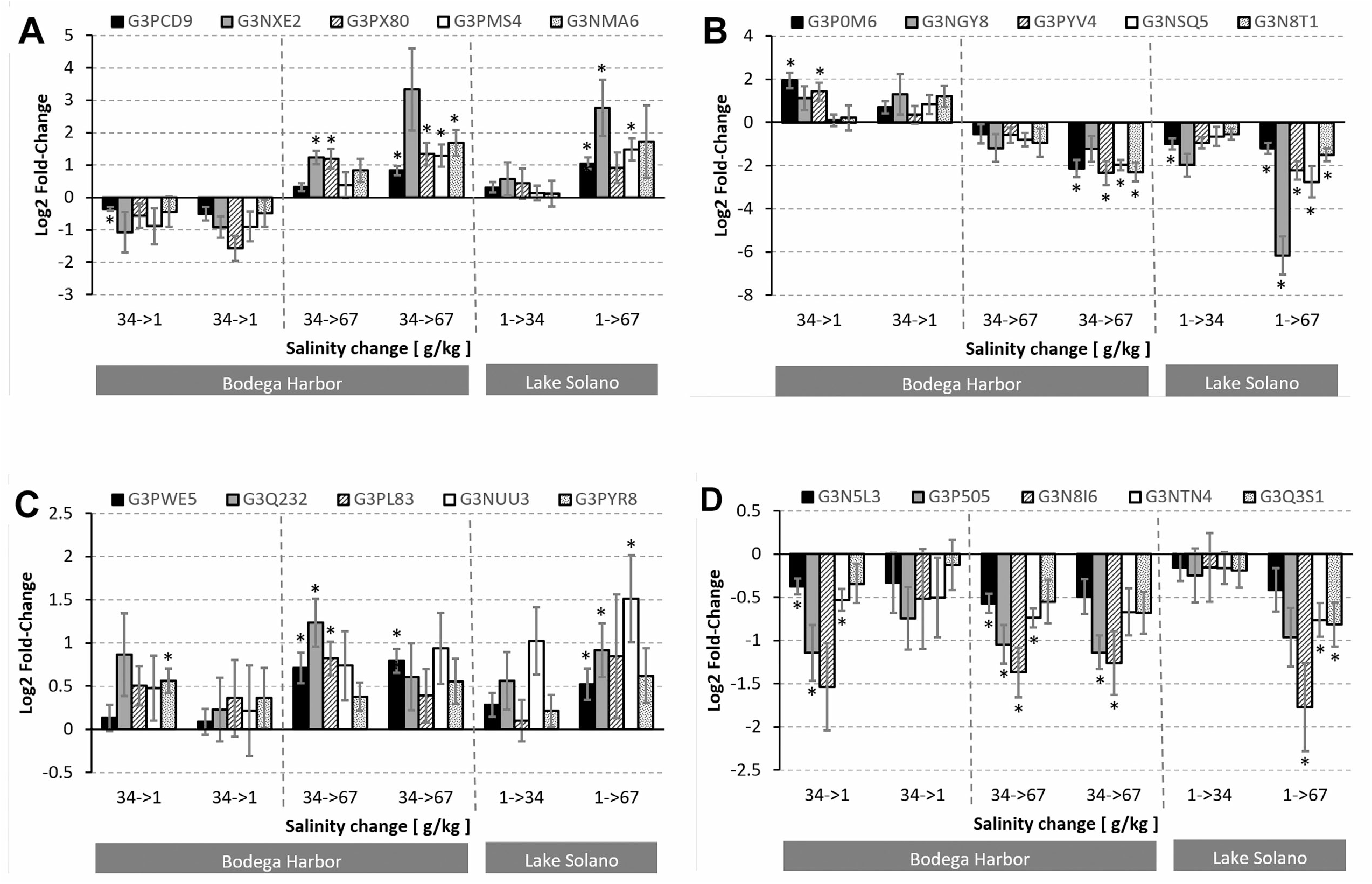
Salinity-induced abundance changes for proteins that correspond to four types of regulatory patterns, which were consistent across all six experiments and significant in at least one experiment. The log2(fold-change) and associated standard error for five exemplary proteins are depicted for each regulatory pattern: **A)** Five proteins whose abundance is directly proportional to salinity (G3PCD9 cytochrome b-c1 complex subunit 2, G3NXE2 aspartate aminotransferase, cytoplasmic, G3PX80 NAD[P] transhydrogenase, G3PMS4 Na^+^/K^+^-ATPase subunit alpha 1, G3NMA6 inositol monophosphatase 1) **B)** Five proteins whose abundance is inversely proportional to salinity (G3P0M6 eosinophil peroxidase, G3NGY8 Golgi-associated plant pathogenesis-related protein 1, G3PYV4 integrin beta-like protein, G3NSQ5 myristoylated alanine-rich C-kinase substrate 1, G3N8T1 anterior gradient protein 2); **C)** Five general stress-induced proteins (G3PWE5 pyruvate kinase, G3Q232 receptor of activated protein kinase C / RACK, G3PL83 grancalcin, G3NUU3 translationally-controlled tumor protein homolog 1, G3PYR8 transaldolase); **D)** Five general stress-repressed proteins (G3N5L3 protein lin-7 homolog B, G3P505 nucleolar transcription factor 1-B, G3N8I6 histone H1, G3NTN4 KHdomain-containing RNA-binding signal transduction-associated protein 1, G3Q3S1 SH3domain-binding glutamic acid-rich-like protein). The Y axis of each graph delineates the six different mesocosm experiments (four for Bodega Harbor fish and two for Lake Solano fish). Asterisks indicates statistically significant abundance changes (multiple-testing corrected P<0.05). Please refer to table S1 for a complete listing of quantitative and statistical data for all proteins that correspond to each of the four regulatory patterns.

### KEGG pathways for osmoregulatory and general stress response proteins

KEGG pathway analysis was used to reveal dominant biological processes associated with each of the four protein regulatory patterns controlled by environmental salinity. The four lists of KO identifiers fitting a protein regulation pattern (proportional to salinity, inversely proportional to salinity, general stress-induced and general stress-repressed) were used with KEGG mapper to identify KEGG pathways that included two or more of the proteins in the corresponding list. The overlap of KEGG pathways associated with the four protein regulation patterns was small and most of the KEGG pathways identified were unique to a single pattern (Figure 6a). However, three KEGG pathways were represented (by different proteins) in both the inversely salinity-regulated and the general stress-repressed protein groups (KEGG 03040 Spliceosome, KEGG 04141 ER protein processing, and KEGG 04810 Regulation of actin cytoskeleton). Two KEGG pathways were shared between proteins that are directly proportional to salinity and general stress-induced proteins (KEGG 00020 Citrate cycle and KEGG 00280 Valine, leucine and isoleucine degradation). The KEGG pathway 04145 (Phagosome) represents all protein regulatory patterns except for the set of proteins whose abundance is directly proportional to salinity. A single KEGG pathway (04915 Estrogen signaling) is represented (by different proteins) in both directly and inversely salinity-regulated proteins. Furthermore, a single KEGG pathway (04510 Focal adhesion) is represented (by different proteins) in both general stress-induced and -repressed proteins. Overlap also exists in a single KEGG pathway (00190 Oxidative phosphorylation) between proteins whose abundance is proportional to salinity and (different) general stress-repressed proteins.

**Figure 6:**
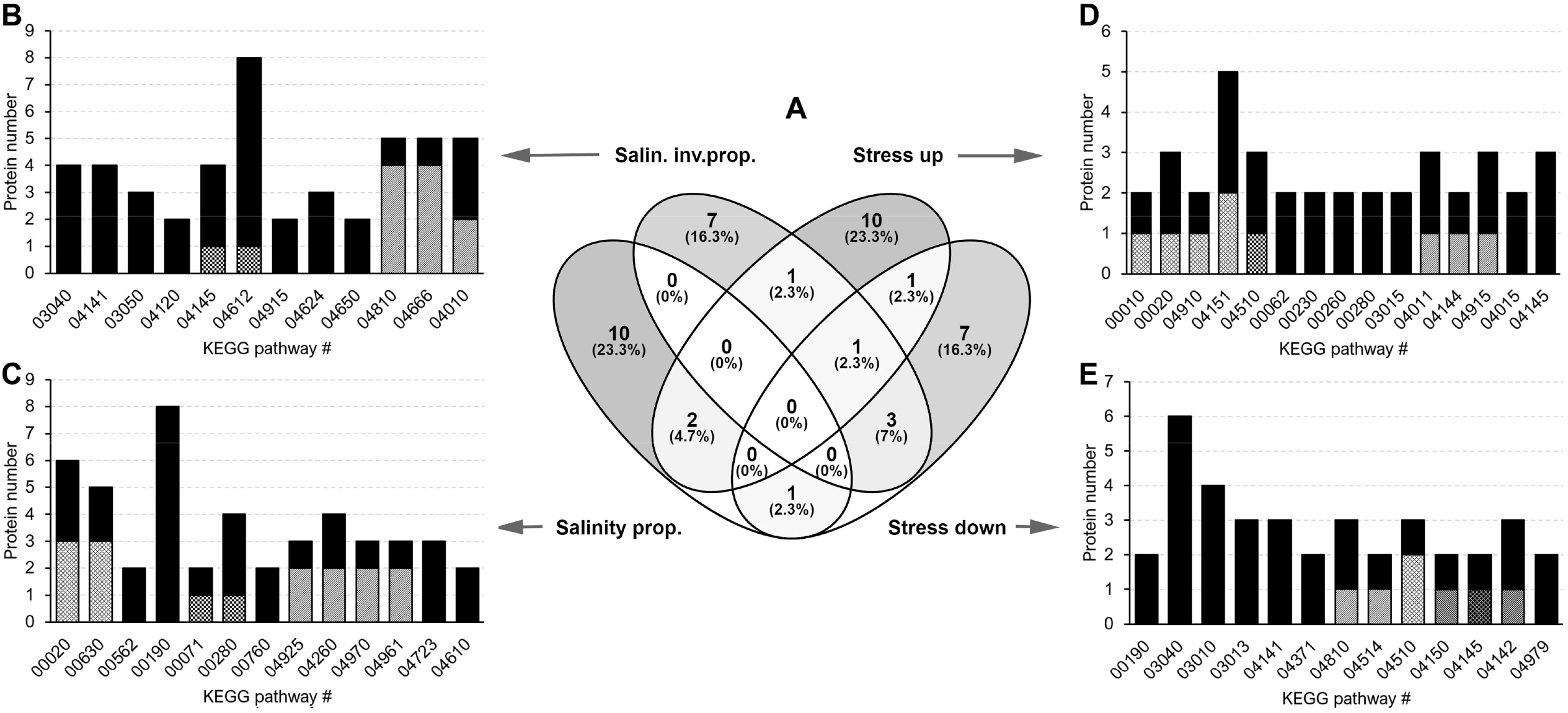
Representation of KEGG pathways fitting four different patterns of salinity-dependent regulation. **A**) Venn diagram illustrating minimal overlap in KEGG pathways associated with the four patterns of protein regulation. The number of proteins that are statistically significant in at least one experiment and display a consistent pattern of regulation in all six experiments is displayed for all matching KEGG pathways (min. 2 proteins/ pathway) in panels B - E. **B)** KEGG pathways for proteins that are inversely proportional to salinity; **C)** KEGG pathways for proteins that are directly proportional to salinity; **D)** KEGG pathways for general stress-induced proteins; **E)** KEGG pathways for general stress-repressed proteins. The shaded area of bars in adjacent KEGG pathways indicates the number of shared proteins that represent more than one pathway. The numbers on the abscissa denote KEGG pathway identifiers. Please refer to the main text or KEGG (https://www.genome.jp/kegg/pathway.html) for assignment of pathway names to these identifiers.

Twelve KEGG pathways consist of at least two proteins that are regulated inversely proportional to salinity. They are KEGG 03040 Spliceosome, 04141 Protein processing in endoplasmic reticulum, 03050 Proteasome, 04120 Ubiquitin mediated proteolysis, 04145 Phagosome, 04612 Antigen processing and presentation, 04915 Estrogen signaling pathway, 04624 Toll and Imd signaling pathway, 04650 Natural killer cell mediated cytotoxicity, 04810 Regulation of actin cytoskeleton, 04666 Fc gamma R-mediated phagocytosis, and 04010 MAPK signaling pathway (Figure 6b). Thirteen KEGG pathways are represented by at least two proteins that are directly proportional in abundance to salinity. These pathways are KEGG 00020 Citrate cycle, 00630 Glyoxylate and dicarboxylate metabolism, 00562 Inositol phosphate metabolism, 00190 Oxidative phosphorylation, 00071 Fatty acid degradation, 00280 Valine, leucine and isoleucine degradation, 00760 Nicotinate and nicotinamide metabolism, 04925 Aldosterone synthesis and secretion, 04260 Cardiac muscle contraction, 04970 Salivary secretion, 04961 Endocrine and other factor-regulated calcium reabsorption, 04723 Retrograde endocannabinoid signaling, and 04610 Complement and coagulation cascades (Figure 6c). The following fifteen KEGG pathways are represented by at least two general stress-induced proteins: 00010 Glycolysis / Gluconeogenesis, 00020 Citrate cycle, 04910 Insulin signaling, 04151 PI3K-Akt signaling, 04510 Focal adhesion, 00062 Fatty acid degradation, 00230 Purine metabolism, 00260 Glycine, serine and threonine metabolism, 00280 Valine, leucine and isoleucine degradation, 03015 mRNA surveillance pathway, 04011 MAPK signaling, 04144 Endocytosis, 04915 Estrogen signaling, 04015 Rap1 signaling pathway, and 04145 Phagosome (Figure 6d). Notably, most of the 34 molecular chaperones/ heat shock proteins (HSPs) included in the DIA assay library did not fit into any of the four regulatory patterns. Only HSP71 (G3NXH8), HSP90beta (G3PJV6), and T-complex1eta (G3Q8L9) displayed the general stress-induced pattern of regulation (Table S1). Thirteen KEGG pathways are represented by at least two general stress-repressed proteins. They are KEGG 00190 Oxidative phosphorylation, 03040 Spliceosome, 03010 Ribosome, 03013 RNA transport, 04141 Protein processing in ER, 04371 Apelin signaling, 04810 Regulation of actin cytoskeleton, 04514 Cell adhesion molecules (CAMs), 04510 Focal adhesion, 04150 mTOR signaling, 04145 Phagosome, 04142 Lysosome, and 04979 Cholesterol metabolism (Figure 6e).

Despite the overlap of some KEGG pathways that contain proteins corresponding to multiple patterns of salinity-dependent regulation, most proteins in these KEGG pathways display a consistent pattern of regulation. For example, even though phosphoenolpyruvate carboxykinase [GTP] (G3NC51) and pyruvate carboxylase (G3Q870) in KEGG pathway 00020 (Citrate cycle) display a general stress-induced pattern of regulation, the abundance of most proteins that constitute this pathway is directly proportional to environmental salinity (Figure 7). Additional examples that illustrate the same trend include KEGG pathways 00280 (Valine, leucine and isoleucine degradation, Figure S1) and 00190 (Oxidative phosphorylation, Figure S2).

**Figure 7:**
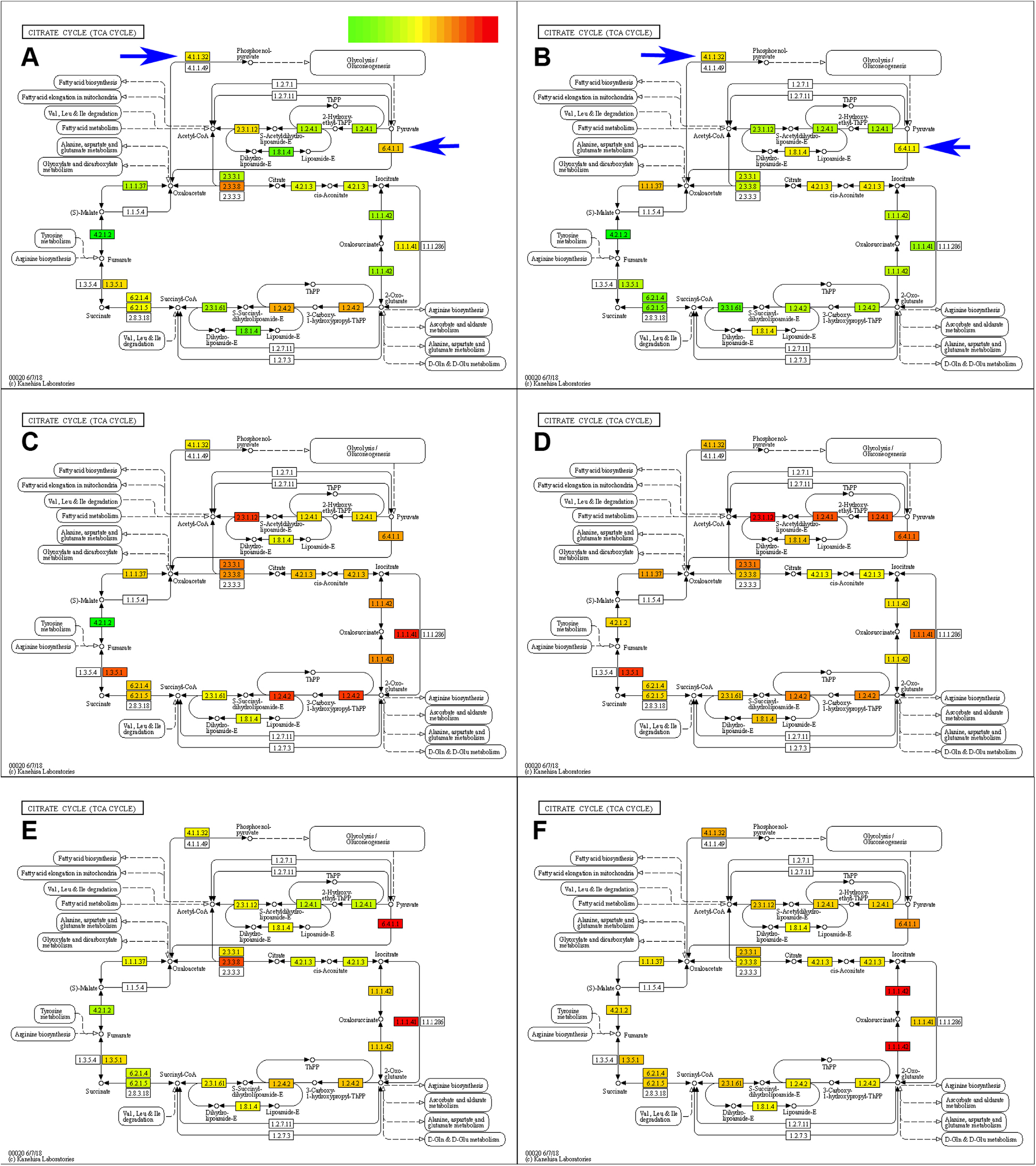
Proportional regulation of KEGG pathway 00020 (citrate cycle) in response to changes in salinity. Proteins that are colored are included in the DIA assay library for *Gasterosteus aculeatus* gill. Color-code represents fold-change (FC) in each experiment according to the legend at the top of panel A (min. green = −4 FC, yellow = no change/ zero FC, max. red = +4 FC). Color-coded FC represents the mean FC relative to the corresponding handling control group for each protein in this pathway in all six mesocosm experiments. **A**) experiment BH1.1 (Bodega Harbor, 34 → 1 g/kg), **B**) experiment BH1.2 (Bodega Harbor, 34 → 1 g/kg), **C**) experiment BH67.1 (Bodega Harbor, 34 → 67 g/kg), **D**) experiment BH67.1 (Bodega Harbor, 34 → 67 g/kg), **E**) experiment LS34 (Bodega Harbor, 34 → 1 g/kg), **F**) experiment LS67 (Bodega Harbor, 34 → 1 g/kg). Blue arrows indicate phosphoenolpyruvate carboxykinase [GTP] (G3NC51) and pyruvate carboxylase (G3Q870), which display general stress-induced regulation patterns that differ from that of many other proteins that are regulated proportionally to salinity (green proteins in panels A and B).

## Discussion

Previously we have revealed pervasive abundance differences in the gill proteome of *G. aculeatus* populations representing different ecotypes (marine, brackish/ anadromous and freshwater) and morphotypes (fully-plated *trachurus* and low-plated *leiurus*) (Li et al. 2018). However, salinity is only one of the environmental parameters that distinguishes habitats in which different populations occur and its contribution to gill proteome differences relative to that of other factors (temperature, food availability, parasitism, etc.) is unknown. The present study was conducted to isolate the effects of salinity on the gill proteome from those of other environmental parameters. This study focuses on the proteome because proteins represent the critical links that determine how genomes interact with the environment to yield specific morphological, physiological, and behavioral phenotypes (Diz et al. 2012; Baer and Millar 2016). On the one hand, proteins are linked directly to specific genes via proteotypic peptides (Keerthikumar and Mathivanan 2017). On the other, proteins represent the critical molecular building blocks that define structure and carry out most biochemical processes and functions of cells, tissues and organisms (Ebhardt et al. 2015). Proteins represent the molecular constituents giving rise to phenotypic variability that is acted upon by natural selection (Clarke 1971, 1971; Frömmel and Holzhütter 1984), which is why almost all targets of pharmaceutical drugs are proteins (Ebhardt et al. 2015). Therefore, analyzing effects of salinity on the proteome provides pertinent insight into molecular mechanisms underlying organismal phenotypes, including salinity tolerance.

Multiple mesocosm experiments were performed on both marine and FW ecotypes and both *trachurus* and *leiurus* morphotypes to permit general conclusions about salinity effects on this species that are independent of eco- or morphotype. In addition, marine fish were acclimated to both FW (1 g/kg) and double-strength SW (67 g/kg) to distinguish general stress response proteins from osmoregulatory proteins. The rationale underlying this distinction is that osmotic homeostasis can by dysregulated in two directions, i.e. the homeostatic range or setpoint can be surpassed during hyperosmotic stress or undercut during hypo-osmotic stress (Kültz 2020). Therefore, specific responses to salinity changes are directional, i.e. osmoregulatory proteins required for hypo-osmoregulation increase proportional to salinity and, conversely, osmoregulatory proteins required for hyper-osmoregulation decrease proportional to salinity. In contrast, general stress response proteins increase irrespective of whether the stress is due to an increase or a decrease of salinity. By filtering out general stress response proteins from the set of gill proteins that change in abundance during salinity acclimation, a more reliable estimate of the contribution of salinity to gill proteome differences observed in field populations can be obtained.

The two populations studied represent typical phenotypes for marine (Bodega Harbor) and landlocked freshwater (Lake Solano) three-spined stickleback ecotypes. Bodega Harbor fish are fully plated while Lake Solano fish are low plated, with a mean lateral plate number of 28 and six, respectively (Knecht et al. 2007). Lake Solano sticklebacks reach a much smaller adult size than Bodega Harbor sticklebacks and this size difference is genetically determined (McPhail 1977; McKinnon and Rundle 2002). The striking morphological differences between Lake Solano FW and Bodega Harbor marine populations are symptomatic of differences between other marine and FW populations of this species (Paepke 1996). Convergence of this phenotype suggests that invasion of low salinity FW habitats harbors a significant cost that drives the fixation of energetically less demanding morphotypes (loss of body armor and dwarfism) based on standing genetic variation. Thus, one would expect differences in the gill proteome of marine and FW sticklebacks, of which there are indeed many as demonstrated by our data (Table S1). However, to maintain focus this study emphasizes conserved osmoregulatory proteins that are most reliable for predicting habitat salinity rather than proteins whose regulation is specific to certain populations, eco-, or morpho-types.

The osmoregulatory proteins that correlate in their abundance with environmental salinity irrespective of eco- or morpho-type include common and previously known ion transporters, e.g. the Na^+^/K^+^-ATPase (NKA) alpha and beta subunits and the Na^+^/K^+^/2Cl^−^ cotransporter (NKCC1). Active transepithelial ion transport is one of the main functions of the teleost fish gill and the activity of the NKA and NKCC1 depends on the osmotic gradient between blood plasma and the environment (Evans 2008). This gradient is more than twice as steep for fish inhabiting SW compared to FW fish, which is reflected in a greater number and larger size of branchial mitochondria-rich ionocytes that produce the necessary ATP (Hiroi and McCormick 2012). In addition, compatible osmolyte synthesizing enzymes (inositol monophosphatase IMPA 1.1 and 1.2), and enzymes involved mitochondrial oxidative metabolism are regulated proportionally to salinity. The latter also reflect an increase in the volume and size of mitochondria-rich ionocytes that parallels the increase in salinity from FW to 2×SW (Conte 2012). Many of these proteins had previously been identified as differently abundant in four *G. aculeatus* populations that occur in habitats ranging from FW to SW (Li et al. 2018). Identification of these proteins is not surprising but establishes salinity as the environmental parameter that underlies their abundance differences in diverse field populations of this species.

Moreover, in addition to these previously known osmoregulatory proteins, our study has identified a novel set of gill proteins and associated functions that differ among field populations (Li et al. 2018) but were not expected to be regulated by salinity. The abundance of two isoforms of Golgi-associated plant pathogenesis protein 1 (GAPR1), interferon-induced protein 44 (INF44), several pattern recognition receptors, and multiple enzymes with known functions in KEGG pathways that regulate immunity and proteostasis processes were negatively correlated with salinity. Currently, studies of GAPR1 in fish other than our previous study (Li et al. 2018) are lacking. However, GAPR1 ortholog abundance is altered during protein aggregation and proliferative diseases that are associated with dysregulation of proteostasis in plants, animals, and humans (Gibbs et al. 2008). Our study shows that the salinity-dependent regulation of several other proteins and KEGG pathways involved in proteostasis mirrors that of GAPR1 proteins, which suggests that the regulation of proteostasis represents a key function of GAPR1 in fish gills. GAPR1 proteins promote tissue remodeling in the mammalian kidney (Baxter et al. 2007) indicating that they could be involved in the remodeling of gill epithelium that is necessary for salinity acclimation of euryhaline fish (Kültz 2015). The salinity-dependent regulation of numerous cell surface antigens (e.g. MHCs), cytokine-inducible proteins (e.g. IFN44, gamma-interferon-inducible lysosomal thiol reductase), and proteins that control cell and epithelial polarity (e.g. anterior gradient protein 2, Cdc42) suggests a primary role in gill epithelium remodeling rather than immunity to adjust the osmoregulatory function of gills. These results are consistent with a previous study of the *G. aculeatus* gill transcriptome, which contains many mRNAs involved in transmembrane ion transport and gill epithelial restructuring that change in abundance during salinity acclimation of both major eco- and morpho-types (Gibbons et al. 2017).

Our data demonstrate that the regulation of these “gill remodeling proteins” does not represent a general stress response. Since these proteins are upregulated during acclimation to hypo-osmotic salinity, their down-regulation during hyperosmotic salinity acclimation cannot be interpreted as an indication of non-specific immunosuppression or stress-induced adjustment of proteostasis. Rather, their regulation is specific to the direction of the osmotic gradient suggesting that they are involved in branchial osmoregulation. Several of these proteins had been identified as up-regulated in gills of FW sticklebacks relative to fish collected from brackish or marine habitats (Li et al. 2018). Their increased abundance in gills of FW fish collected from Lake Solano had previously been interpreted as an indication of either parasitism, methylmercury toxicity, or FW specific pathogens that induce immune defenses. However, the present study clearly shows that salinity alone can account for the altered abundance of these proteins and that they are regulated in the same way in marine Bodega Harbor fish as in FW Lake Solano fish.

In summary, this study utilizes DIA quantitative proteomics to identify proteins involved in osmoregulation and general stress response proteins whose abundance changes in response to salinity acclimation of *G. aculeatus*. Mapping these protein sets to corresponding KEGG pathways shows that proteins involved in osmoregulation are associated with KEGG pathways that differ from those associated with non-specific stress response proteins. Gill proteins that are directly proportional to salinity function in transepithelial ion transport, compatible osmolyte synthesis and mitochondrial oxidative metabolism. Gill proteins that are inversely proportional to salinity exert their osmoregulatory function by controlling tissue remodeling, cell polarity, and proteostasis processes. The corresponding KEGG pathways provide insight into key molecular mechanisms that are adjusted in *G. aculeatus* gills when salinity changes. Moreover, our results show that salinity sufficiently explains the differential abundance of these osmoregulatory proteins in populations inhabiting FW and SW.

---

**Figure S1:** Salinity-dependent regulation of KEGG pathway 00280 (Valine, leucine and isoleucine degradation). Proteins that are colored are included in the DIA assay library for *Gasterosteus aculeatus* gill. Color-code represents fold-change (FC) in each experiment according to the legend at the top of panel A (min. green = −4 FC, yellow = no change/ zero FC, max. red = +4 FC). Color-coded FC represents the mean FC relative to the corresponding handling control group for each protein in this pathway in all six mesocosm experiments. **A**) experiment BH1.1 (Bodega Harbor, 34 → 1 g/kg), **B**) experiment BH1.2 (Bodega Harbor, 34 → 1 g/kg), **C**) experiment BH67.1 (Bodega Harbor, 34 → 67 g/kg), **D**) experiment BH67.1 (Bodega Harbor, 34 → 67 g/kg), **E**) experiment LS34 (Bodega Harbor, 34 → 1 g/kg), **F**) experiment LS67 (Bodega Harbor, 34 → 1 g/kg).

**Figure S2:** Salinity-dependent regulation of KEGG pathway 00190 (Oxidative phosphorylation). Proteins that are colored are included in the DIA assay library for *Gasterosteus aculeatus* gill. Color-code represents fold-change (FC) in each experiment according to the legend at the top of panel A (min. green = −4 FC, yellow = no change/ zero FC, max. red = +4 FC). Color-coded FC represents the mean FC relative to the corresponding handling control group for each protein in this pathway in all six mesocosm experiments. **A**) experiment BH1.1 (Bodega Harbor, 34 → 1 g/kg), **B**) experiment BH1.2 (Bodega Harbor, 34 → 1 g/kg), **C**) experiment BH67.1 (Bodega Harbor, 34 → 67 g/kg), **D**) experiment BH67.1 (Bodega Harbor, 34 → 67 g/kg), **E**) experiment LS34 (Bodega Harbor, 34 → 1 g/kg), **F**) experiment LS67 (Bodega Harbor, 34 → 1 g/kg).

