## Supplementary figures and images for "Proteomics of osmoregulatory responses in threespine stickleback gills"

### Supplemental Figure 1

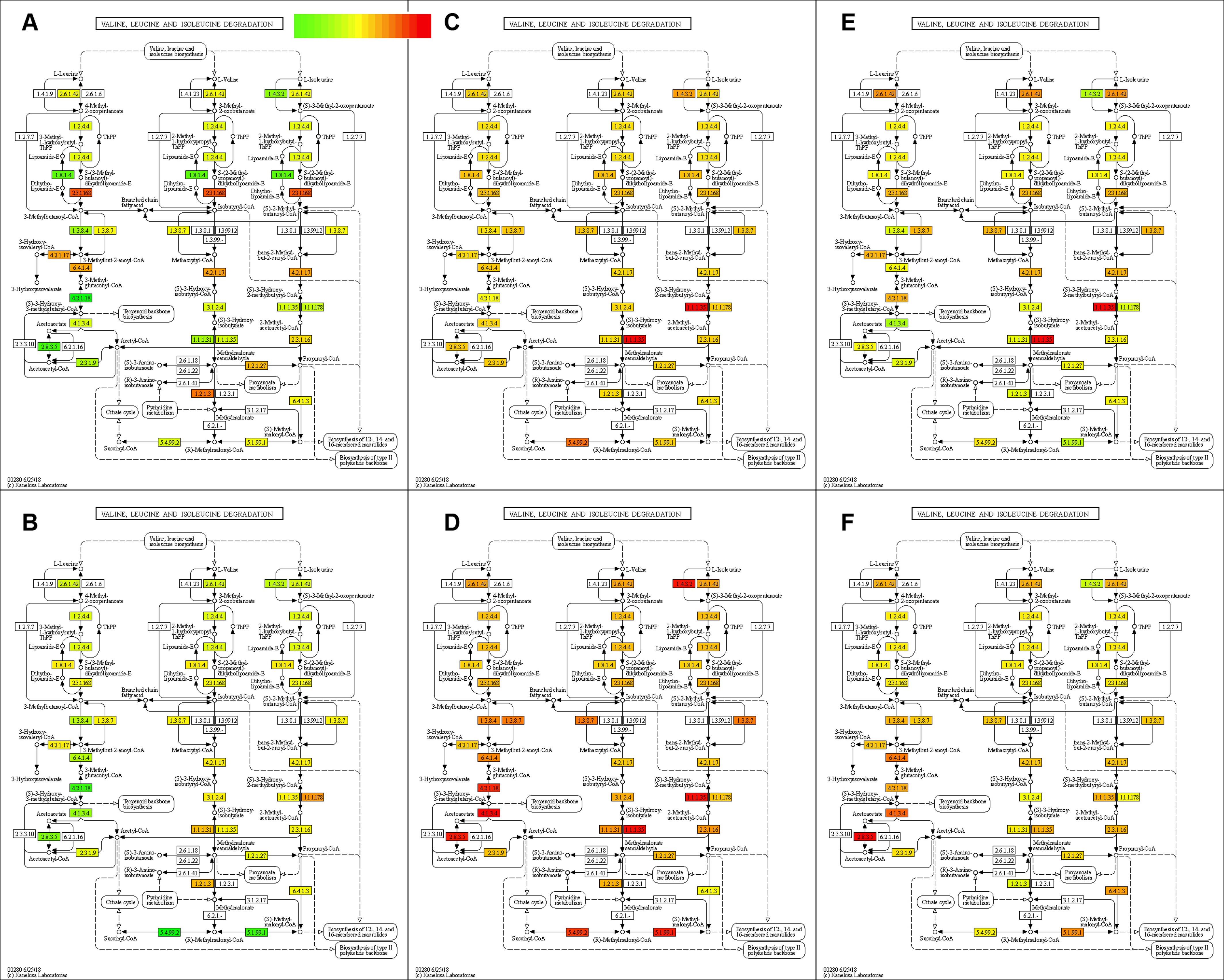

### Supplemental Figure 2

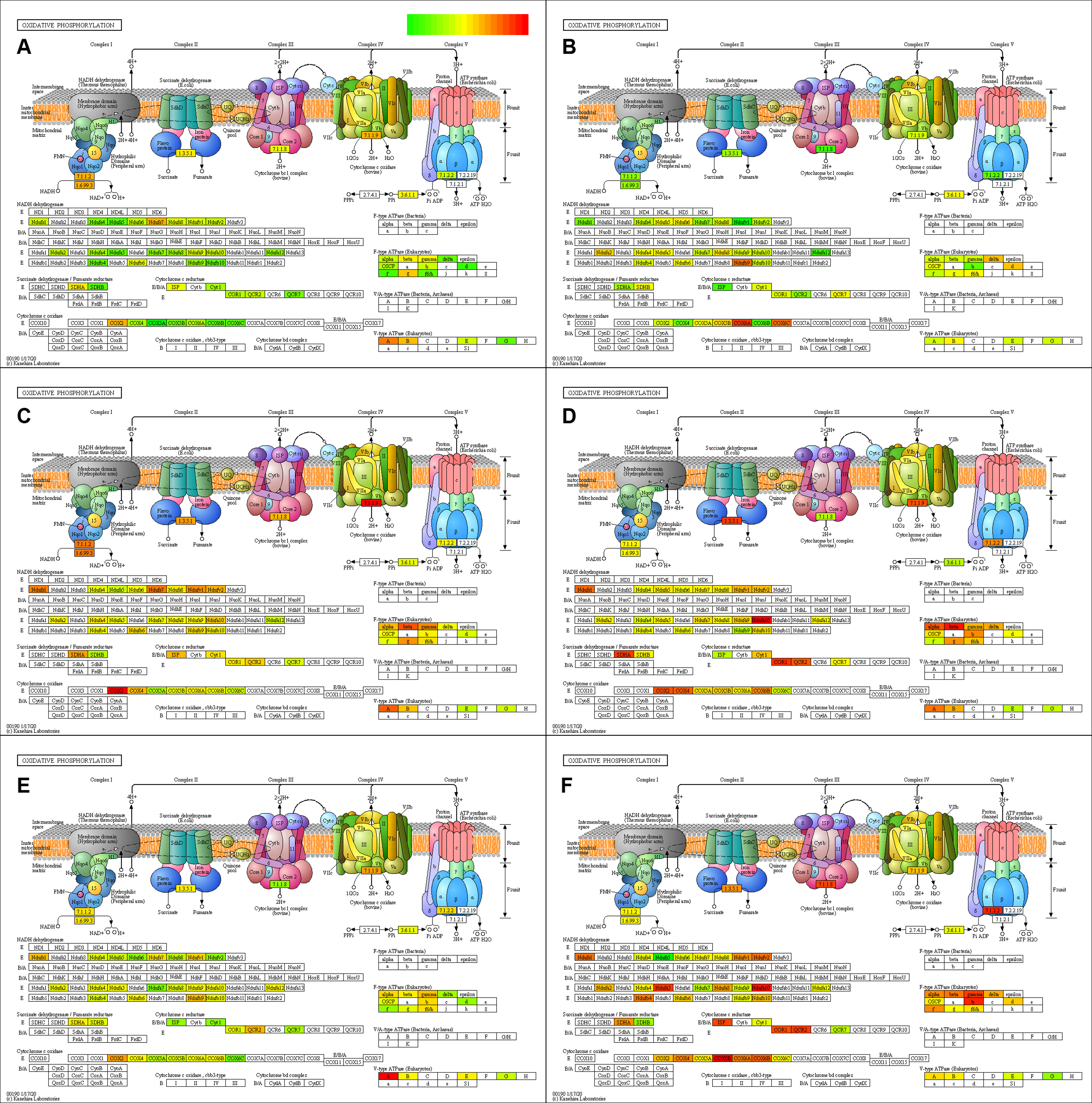
